# Lis1 regulates Phagocytosis by co-localising with actin in the Phagocytic cup

**DOI:** 10.1101/835769

**Authors:** Aditya Chhatre, Paulomi Sanghavi, Roop Mallik

## Abstract

Phagocytosis, the ingestion of solid particles by cells, is essential for nutrient uptake, innate immune response, antigen presentation and organelle homeostasis. Here we show that Lissencephaly-1 (Lis1), a well-known regulator of the microtubule motor Dynein, also co-localises with actin at the phagocytic cup in the early stages of phagocytosis. Both knockdown and overexpression of Lis1 perturbs phagocytosis, suggesting that an optimum level of Lis1 is required to regulate actin dynamics within the phagocytic cup during particle engulfment. This requirement of Lis1 is replicated in mouse macrophage cells as well as in the amoeba *Dictyostelium*, indicating an evolutionarily conserved role for Lis1 in phagocytosis. In support of these findings, a general role for Lis1 in regulating actin dynamics is suggested by observing defective migration of cells overexpressing Lis1. Taken together, Lis1 localises to the phagocytic cup and influences actin dynamics in a manner that is important for the uptake of solid particles in cells.

## Introduction

Phagocytosis involves ingestion of foreign particles larger than ~500nm by a cell. While lower protozoic heterotrophs rely on phagocytosis for nutrition, specialized cell types in higher organisms employ phagocytosis for innate immune response, antigen presentation and clearance of dead tissue. Although the purpose of phagocytosis might vary between cell types and organisms, the fundamental steps governing phagocytosis remain conserved through evolution. Phagocytosis is strongly dependent on the actin cytoskeleton, wherein actin acts as a scaffold to form membrane protrusions around the particle at the cell cortex (reviewed in [1]). Actin is subject to regulation by a host of actin associated proteins including nucleating, filament severing, end-capping, bundling, membrane tethering, and scaffolding proteins. Lissencephaly-1 (Lis1) is a protein that is classically known to interact with, and modulate the function of microtubule-based dynein motors [2][3]. Mutations in *LIS1* cause severe developmental defects in formation of the brain (Lissencephaly). Mutations in dynein mirror the defects in neuronal migration caused by Lis1, prompting much interest on a common disease-relevant genetic pathway involving Lis1, dynein and microtubules[4].

However, outside this classical dynein-dependent pathway, Lis1 also acts as a scaffold for actin-nucleating machinery at the leading edge of neuronal growth cones in the developing brain [5]. Lis1 also cross-talks with the extracellular adhesion complex at the focal adhesions of fibroblasts [6], and is necessary for generating forces against the substratum for cell migration. Knockdown of Lis1 reduces F-actin concentration at the leading edge of neurons [7]. These observations suggest an additional role for Lis1 in actin-dependent remodeling of membranous structures. Actin driven remodeling of the plasma membrane also happens, albeit on a localized scale, during phagocytosis. Actin nucleation at the site of phagocytosis generates the force to form protrusions (phagocytic cup) around the particle. Upon completion of engulfment, the particle is enclosed in an actin containing lipid-protein bilayer membrane to become a phagosome, which then goes through a programmed maturation pathway towards lysosomal fusion and degradation [8]. Therefore, both phagocytosis and cell migration involve extracellular adhesion complexes, actin cytoskeleton and deformations in the plasma membrane.

Keeping in mind the above similarities, we investigated a potential role for Lis1 in regulating phagocytosis in mouse macrophage cells, and also in the highly phagocytic social amoeba *Dictyostelium*. Lis1 is a member of the conserved WD (tryptophan-aspartic acid) family of proteins [3][4] that have multiple C-Terminal domains for interaction with other proteins. A role for Lis1 in controlling actin dynamics was revealed in *Dictyostelium* [11]. *Dictyostelium* Lis1 (DdLis1) and human Lis1 are 53% identical and have similar domain structure. It was proposed that impairment of DdLis1 reduces F-actin content to alter actin dynamics in cells, and a direct interaction of DdLis1 with DdRac1 (a Rho-GTPase) was also shown, suggesting that DdLis1 promotes actin polymerization via the well-known Rho-GTPase pathway [11]. When DdLis1 function was impaired, dynein mediated delivery of DdRac1 to the cell cortex was blocked, suggesting an alternate (dynein-dependent) mechanism for inhibiting actin dynamics resulting from the lack of DdRac1 delivery to the cell cortex via dynein. These results highlight Lis1 as an important regulator of the actin machinery via a direct actin-dependent mechanism, or an indirect dynein dependent pathway.

Here we use knockdown and overexpression studies to show that Lis1 co-localises with actin in the phagocytic cup. Perturbations in Lis1 lead to defects in phagocytosis including reduced phagocytic efficiency, incomplete formation of the phagosomal cup and reduced binding of particles to cell surface. Such perturbations also lead to defects in cell motility, suggesting that Lis1 alters actin dynamics to cause defects in phagocytosis. Lis1, along with actin-associated proteins, is enriched on phagosomes at an early stage but is lost as the phagosome undergoes maturation inside cells. Taken together, a specific role for Lis1 is suggested in actin-dependent remodelling of membranes around particles during early stages of the phagocytic pathway.

## Results

### Effects of Lis1 Knockdown and Overexpression on Phagocytosis

To assay for phagocytic efficiency we fed silica beads (*D* = 2.5μm diameter) to RAW 264.7 macrophages that had been transfected with either scrambled shRNA or Lis1 specific shRNA using an electroporation technique [12]. To identify the cells that had been transfected, a GFP plasmid was co-transfected with the Lis1 shRNA plasmids. Supplementary Fig 1A verifies the knockdown of Lis1 in RAW macrophages. Beads were incubated with RAW 264.7 macrophages at a density of 100 beads/cell for 5mins. Cells were then fixed and imaged on a confocal microscope. Beads appeared as dark hollow circles on a green background in such images (Fig 1A, left panel), allowing us to easily count the number of beads in individual cells. Fig 1A shows single representative confocal sections of RAW cells. To count the total number of beads/cell (right panel, Fig 1A), we summed up all beads in Z-stacks across the depth of a cell, making sure not to count a given bead more than once. The mean number of beads/cell was taken as a measure of phagocytic efficiency. shRNA-mediated knockdown of Lis1 resulted in loss of phagocytic efficiency compared to scrambled shRNA (Fig 1A; right panel), suggesting that Lis1 is required for efficient phagocytosis. Next, to investigate the effect of increasing Lis1 beyond normal levels, we overexpressed GFP-Lis1 in these cells using the same electroporation protocol used for shRNA transfection. GFP cDNA was transfected as a control. Because Lis1 knockdown inhibited phagocytosis (Fig 1A), we expected to observe enhanced phagocytosis after Lis1 overexpression. Contrary to our expectations, and similar to Lis1 knock-down, overexpression of GFP-Lis1 also reduced bead uptake (Fig 1B). As discussed earlier Fig 1B (left panel) shows a single Z-section, but the total number of beads/cell (Fig 1B, right panel) was obtained after counting beads across all Z-sections.

**Fig 1.**
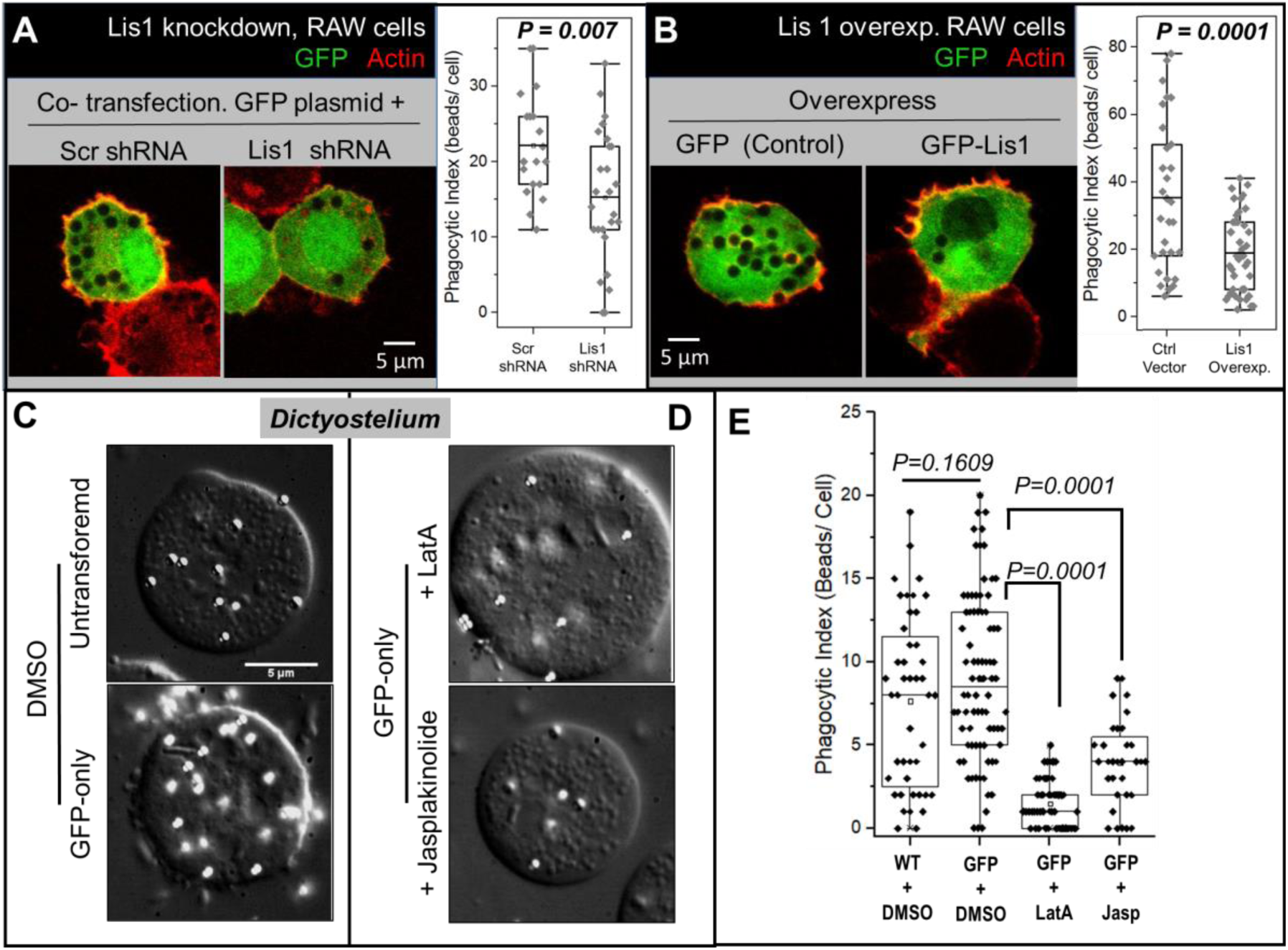
Optimal levels of Lis1 are required for early events in phagocytosis. Similarly, Optimal Actin balance is necessary for phagocytosis. A. A single confocal section of RAW 264.7 macrophage cells co-expressing GFP plasmid with either scrambled shRNA or Lis1 shRNA. Scatter dot plot (right panel) shows quantifications of no. of beads per cell across all the z-stacks imaged. Cells were incubated at a ratio of 100 beads per cell and counterstained with Rhodamine phalloidin (See methods). B. A single confocal section of RAW 264.7 macrophage cells expressing either control GFP or GFP-Lis1 plasmids. Scatter dot plot (right panel) shows quantifications of no. of beads per cell across all the z-stacks imaged. C. DIC images of either WT or GFP overexpressing *Dictyostelium* cells treated with 750nm polystyrene beads (ratio of 100 beads per cell) in presence of DMSO. Polystyrene beads appear as bright circles of uniform diameter. Scale Bar = 5 microns in all cases. D. DIC images of Dictyostelium AX2 cells stably overexpressing control GFP construct treated with 750nm polystyrene beads similar to C, in presence of Jasplakinolide or LatrunculinA as indicated. E. Scatter dot plots showing phagocytic index of cells of each type as mentioned in C and D.

### Effects of Actin stabilization and depolymerization on Phagocytosis

The above observations suggest that an optimum level of Lis1 is required, possibly to regulate actin dynamics, during phagocytic uptake of particles. To investigate further we used pharmacological agents that destabilize or stabilize F-actin in cells. *Dictyostelium* cells overexpressing GFP-only (control) exhibited robust phagocytosis comparable to untransfected cells (Fig 1C, 1E). However, bead uptake was inhibited when cells were treated with 0.3µM LatrunculinA (LatA; Fig 1D, 1E). LatA sequesters cellular G-actin leading to depolymerisation of actin filaments [13]. We next treated cells with Jasplakinolide, a drug that induces actin polymerization or stabilises pre-existing F-actin filaments by directly binding to them [14]. We again observed inhibition of phagocytosis when cells were treated with Jasplakinolide (Fig 1D, 1E). Thus, an optimal balance of F and G actin appears necessary in cells for proper phagocytic cup formation. Our finding agrees with the results of genetic perturbation where the actin cytoskeleton was found necessary for particle uptake by phagocytosis [15] [1]. Stabilizing, as well as destabilizing F-actin cytoskeleton by genetically disrupting its modulators was found to cause defects in phagocytic uptake[16][17].

It was shown that expression of a dominant negative mutant of Lis1 reduces F-actin concentration at the leading edge of neurons [7]. Maltose binging protein (MBP) tagged Lis1 overexpressed in *Dictyostelium* was also inferred to act in a dominant negative manner, with the effect that radially organized microtubules were disrupted and the nucleus-to-centrosome distance was increased[11]. In contrast to that study, we did not find any gross defect in the organization of microtubules in His-GFP-Lis1 overexpressing cells (Fig 2A), and the distance between nucleus and centrosome was also unchanged (Fig 2B). This suggests that the GFP-Lis1 overexpression construct used in our studies functions like endogenous (normal) Lis1, and does not cause a dominant negative effect like the MBP tag [11]. This was further confirmed when His-GFP-Lis1 could pull down endogenous Dynein from *Dictyostelium* cell lysate (Fig 2C), suggesting that it is functioning “normally” (i.e. it can interact with dynein). Because Lis1 +/− heterozygous neurons show a reduction in F-actin content [5], it is possible that endogenous Lis1 promotes formation of F-actin in cells. The overexpressed His-GFP-Lis1, (which presumably functions like endogenous Lis1) could therefore increase the amount of F-actin locally at the actin cup, and this excess actin could inhibit the dynamic re-organization of actin that is required for cup formation around phagosomes. To explore this possibility, we treated GFP-Lis1 cells with 0.3µM LatA in order to destabilize F-actin and rescue actin dynamics towards “normal” levels. Indeed, this treatment led to a mild rescue of the phagocytic index (Figs 2D, 2E). However, we did not observe any obvious difference in the bulk F/G actin ratio in an actin pelleting assay (data not shown), suggesting that the effect of Lis1 on actin dynamics is local, and kicks in only when the cup has to be formed.

**Fig 2.**
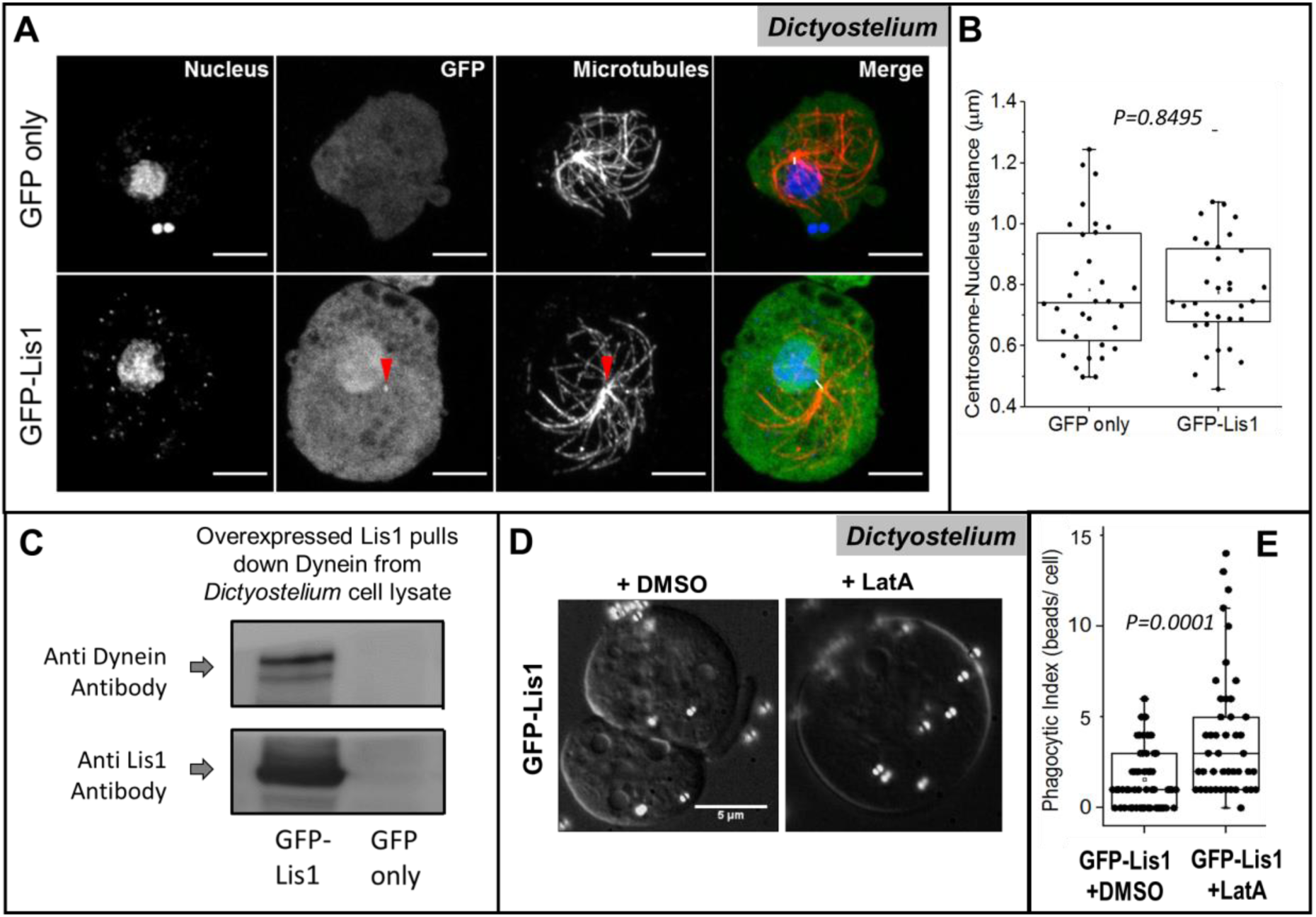
Microtubule organization and centrosome-nucleus distance is not perturbed upon Lis1 overexpression in Dictyostelium cells. Actin depolymerisation rescues the phagocytic defect in Lis1 overexpressing cells. A. Dictyostelium cells overexpressing either GFP or GFP-Lis1 constructs. Radial organization of microtubules is maintained in both the cases. Images are maximum intensity projections of z-stacks taken through the thickness of cells. Notice that a bright green spot is observed at the centrosome (red arrowhead) in GFP-Lis1 case but not for GFP-only case. The centrosomal localisation of Lis1 is in agreement with report in[11]. Scale Bar = 5μm. B. Box plots showing the distance between centrosomes and nuclei of cells. n = 32 cells in each case., Student’s t-test. C. Pull down of Dictyosteium cell lysates using Nickel NTA beads to detect interaction of overexpressed Lis1 with endogenous dynein. Cells used were overexpressing His-GFP-Lis1 or His-GFP (negative control). Dynein heavy chain is detected only in pulldown fraction of His-GFP-Lis1. Lis1 antiserum was used to detect the Lis1 protein. Equal protein amount was used for both pulldown samples. D. DIC images of Dictyotelium AX2 cells stably overexpressing GFP-Lis1 construct treated with 750nm polysterene beads (100 beads per cell) in presence of DMSO or Latrunculin. Scale Bar=5µm. E. Scatter dot plots showing phagocytic index of cells of each type as mentioned in C. Student’s T-test. n = 30-45 cells across 2-3 independent experiments.

### Perturbations in Lis1 cause incomplete phagocytic cup formation

The above results suggest that Lis1 influences actin dynamics, and an optimal level of Lis1 is required for sustaining actin-driven dynamics during phagosomal cup formation. Perturbation in Lis1 level may lead to inefficient phagocytosis, perhaps by restricting actin cup formation around phagosomes. To test this, we next asked whether phagocytic cup formation is affected upon Lis1 expression. *Dictyostelium* cells can be pulsed with beads for a short period (5 minutes) to prepare “early phagosomes”, or pulsed followed by a chase period (40 minutes) to prepare “late phagosomes” as described by us [18][19]. This allowed us to assay for actin dynamics at a defined (early) stage of phagocytic uptake in *Dicytostelium*. Rhodamine-phalloidin was used to visualize actin in the phagocytic cup around phagosomes[15]. Formation of actin around a bead is shown schematically in Fig 3A. The use of beads of defined size (diameter *D* = 2.5μm) allowed us to quantify the extent of cup formation around a single bead. To do this we measured the arc length of the actin cup (= ***d***; see Fig 3A) around phagocytosed beads, and then calculated the fraction of total bead circumference (=π*D*) that was covered by the actin cup. Fig 3B shows images of phagocytosed beads in cells. Actin is stained using rhodamine-phalloidin and beads are visualized using differential interference contrast (DIC) microscopy. The insets in Fig 3B show the actin cup around individual phagosomes (images in inset were inverted for clarity). The arc was linearized as a function of the angle (θ) around a circle centred at the bead by converting the image into polar coordinates (Fig 3C). The line intensity of this linearized arc was plotted as a function of θ (Fig 3D). We then measured the % of the circumference covered by the actin cup for individual beads [=100-(*d*/π*D*\*100)]. We observed a significant reduction in phagocytic cup engulfment after Lis1 overexpression (Fig 3E), showing that cup formation is defective and/or delayed in Lis1 overexpressing cells. Attempts to prepare Lis1 knockdown *Dictyostelium* cells were not successful, perhaps because it is a haploid organism causing Lis1 knockdown to be lethal.

**Fig 3.**
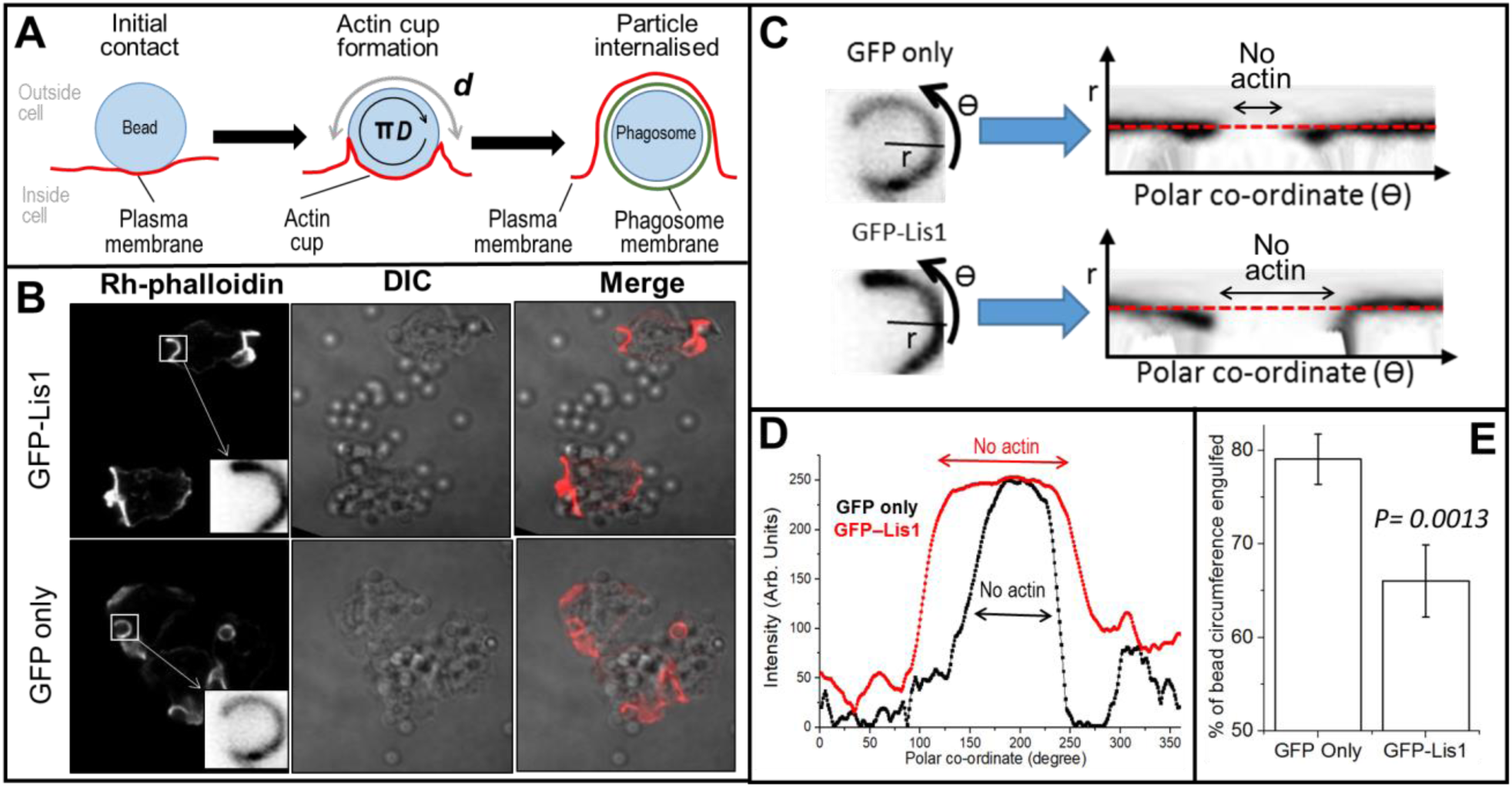
Overexpression of Lis1 causes defect in phagocytic cup progression. A. Simplified schematic showing progression of Actin rich phagocytic cup leading to engulfment of bead. d = length of arc un-engulfed at given time-point t; D = diameter of bead. %bead engulfed = 100-(d X 100/πD). When d = 0, bead is completely engulfed. B. Representative confocal images of Dictyostelium (AX2) cells either expressing GFP-only (control) or Lis1-GFP plasmids, counterstained with rhodamine-phalloidin. Inset: LUT inverted hi-mag images of partially engulfed 2.5um silica beads. Cells were fixed and stained at timepoint t = 125sec post pulse. C. ROI of either GFP only or GFP-Lis1 cups are shown along with their polar transforms. Rhodamine labelled arcs (inverted LUTs) are represented by line segment when radial segment (r) sweeps the angular function (Ɵ). D. Line intensity plots of red dotted lines shown in A. Notice the shorter FWHM for GFP-only cells. E. Bar graphs showing lengths of arcs (as % of total bead circumference) engulfed by cell. n = 45 beads across 3 independent experiments in each case.

### Lis1 localizes to the Actin cup on Phagosomes

Lis1 has been shown to interact physically with actin in-vitro[5][11]. Since we observed that Lis1 modulates actin dependent processes during early phagocytosis, we hypothesized that Lis1 might be localized at the site of phagocytosis itself. To verify this, we imaged GFP in RAW cells transfected with Lis1-GFP or GFP (control). Actin was imaged using rhodamine-phalloidin (Fig 4A). The intensity of GFP and actin were measured along a radially directed line starting at the centre of phagocytosed beads and progressing into the interior of the cell beyond the bead radius (see Fig 4A inset). The averaged intensity profile across multiple phagosomes obtained in this manner is shown in Fig 4B. The profile for Lis1-GFP intensity matched closely with actin, both peaking at ~1.3μm (≈ bead radius) from the centre of phagosomes. In contrast, no obvious peak was observed in control (GFP-only) cells, with GFP intensity remaining fairly elevated beyond the radius of the phagosome. This suggests that Lis1 co-localises with actin at the actin cup around phagosomes at an early stage. Note that absolute intensity levels of GFP signal cannot be compared between these two overexpression conditions because expression levels of GFP and GFP-Lis1 may be different.

**Fig 4.**
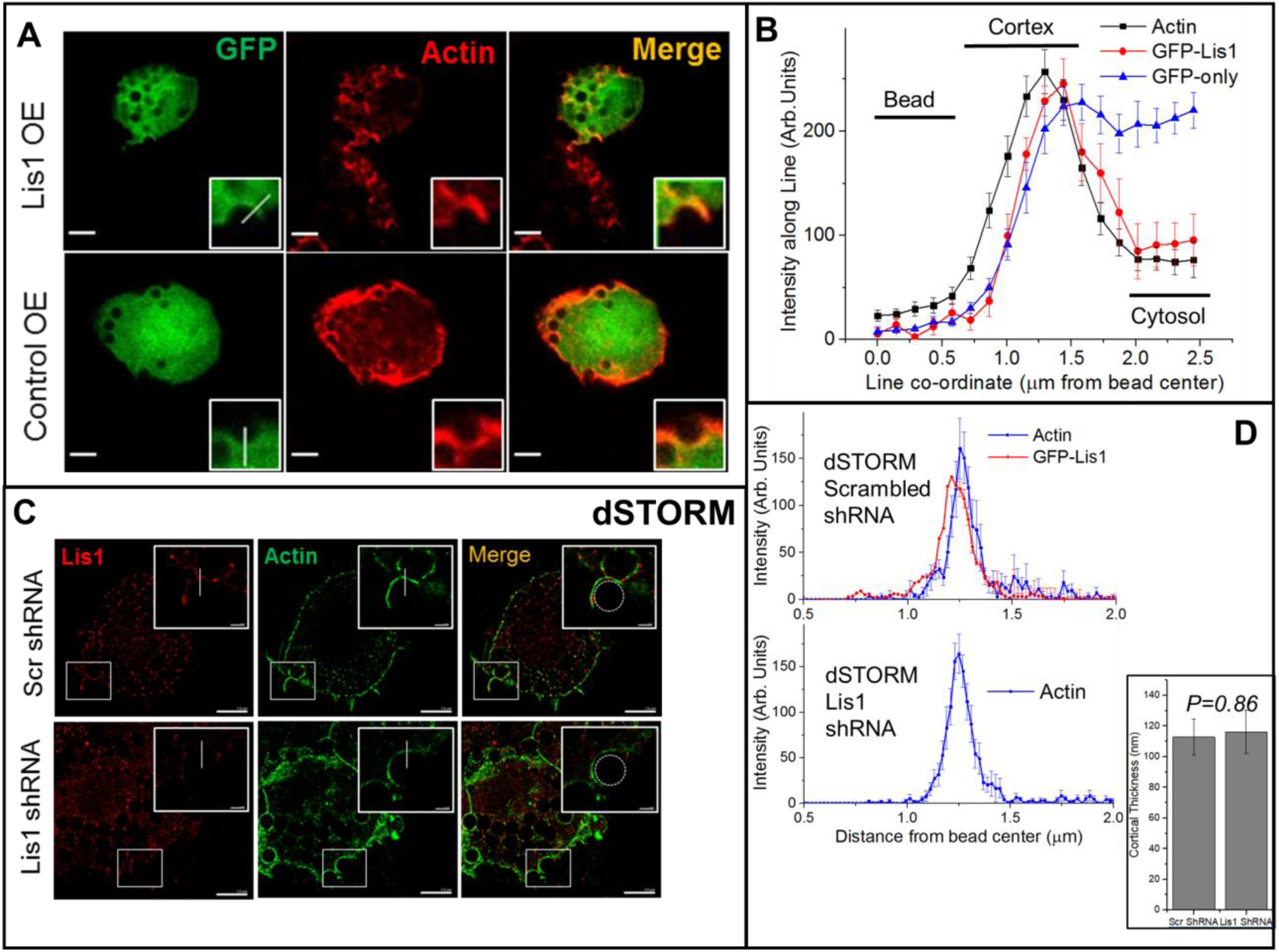
Lis1 localizes to the phagocytic cup. A. Representative confocal images of RAW cells overexpressing either GFP-Lis1 or control GFP constructs, fed with 2.5um silica beads. Insets: Hi-mag of phagocytic cups. Scale bars = 5µm. B. Intensity traces along the indicated Line ROIs in phagocytic cups. Note the relative enrichment of Lis1-GFP at the cortex (marked by Actin trace) as compared to Ctrl-GFP, which is uniform in the cytosol. Error bars are SEM. n = 6-15 beads in 3 independent experiments. C. Representative dSTORM images of RAW cells on scrambled shRNA or Lis1 shRNA background, fed with 2.5um silica beads. Scale bars = 5μm. Insets: Hi-mag of phagocytic cups with Line ROIs. Scale bars = 1um. D. Intensity traces along indicated Line ROIs in phagocytic cups. Note the enrichment of Lis1 at the cortex marked by Actin trace in scr shRNA, which is absent in Lis1 shRNA. Error bars are SEM. n = 7 beads in 2 independent experiments. Inset: Average (±SEM) thickness of cortex (measured by FWHM of Actin plots in D) for mentioned conditions.

To verify that endogenously present Lis1 co-localises precisely with the actin cup in cells, we performed dSTORM super-resolution imaging of phagocytic cups in RAW cells (Methods). Endogenous Lis1 was imaged using anti-Lis1 antibody and a secondary antibody labelled with Alexa-555. Alexa 488-phalloidin was used to image actin. Lis1 was indeed enriched at the phagocytic cup, where it co-localized with actin (Fig 4C). The inset in Fig 4C shows a radially directed line starting at the centre of a phagosome, and Fig 4D (upper panel) shows the fluorescence intensity along such lines averaged across multiple phagosomes. Alexa-555 (marks Lis1) and Alexa-488 (marks actin) both showed a sharp peak exactly at 1.25 μm (= bead radius). The peak in actin was closely coincident with a similar peak for Lis1, suggesting that both proteins localize to the surface of the early phagosome. The width of this peak was 112.8±11.7 nm, in agreement with reports of the thickness of cortical actin [20][21]. Fig 4D (lower panel) shows the averaged peak for actin observed in Lis1 shRNA knockdown cells. The peak for actin has a width (=116.2±14.2 nm), statistically same as the width in control cells (*P* = 0.86). These findings suggest that there is no gross perturbation in the thickness of the actin cup surrounding the phagosome after Lis1 knockdown. Rather, the (lateral) dynamics of actin on the phagosome surface may be perturbed because the actin cups are unable to extend around the phagosome in absence of Lis1. We also note that localization of Lis1 to the cell cortex was not perturbed in Lis1 overexpressing cells (compare Figs 1A, 1B).

### Perturbed chemotaxis upon Lis1 overexpression

To investigate if Lis1 overexpression causes perturbation in actin dynamics, we employed a chemotaxis assay to measure cell migration of *Dictyostelium*. *In-vitro* cell migration assays have been used to analyse the effect of perturbations on the cytoskeleton [3][4][23], including Lis1 perturbation [7], where it was observed that radial migration of neuronal cells (grown as re-aggregate cultures) was grossly reduced upon Lis1 knockdown. For the purpose of this study, we used an under-agarose chemotaxis protocol for *Dictyostelium* adapted from[24](Methods). Briefly, a concentration gradient of chemo-attractant source is prepared in agar troughs in petri plates. Subsequently, the cells are exposed to this gradient and their migration recorded by video microscopy.

Upon subjecting GFP overexpressing cells (control) to a folate gradient, the cells exhibited robust motion (Fig 5A). Individual cells were tracked using the manual tracker plugin of ImageJ. The migration speed was 11.6±0.55 µm/min for control (GFP-only) cells. These cells moved persistently in directed manner up along the folate gradient. As compared to controls, GFP-Lis1 overexpressing cells showed slower migration (speed = 5.34±0.33 µm/min; Fig 5B). The persistence of motion during a defined time interval was estimated from the tortuosity (***τ***). The meaning of ***τ*** is illustrated in the inset of Fig 5B for a period of motion starting at (x_1_,y_1_) and ending at (x_2_,y_2_).

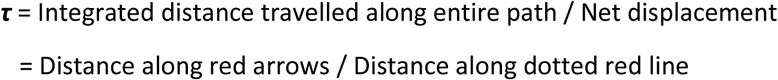

**Fig 5.**
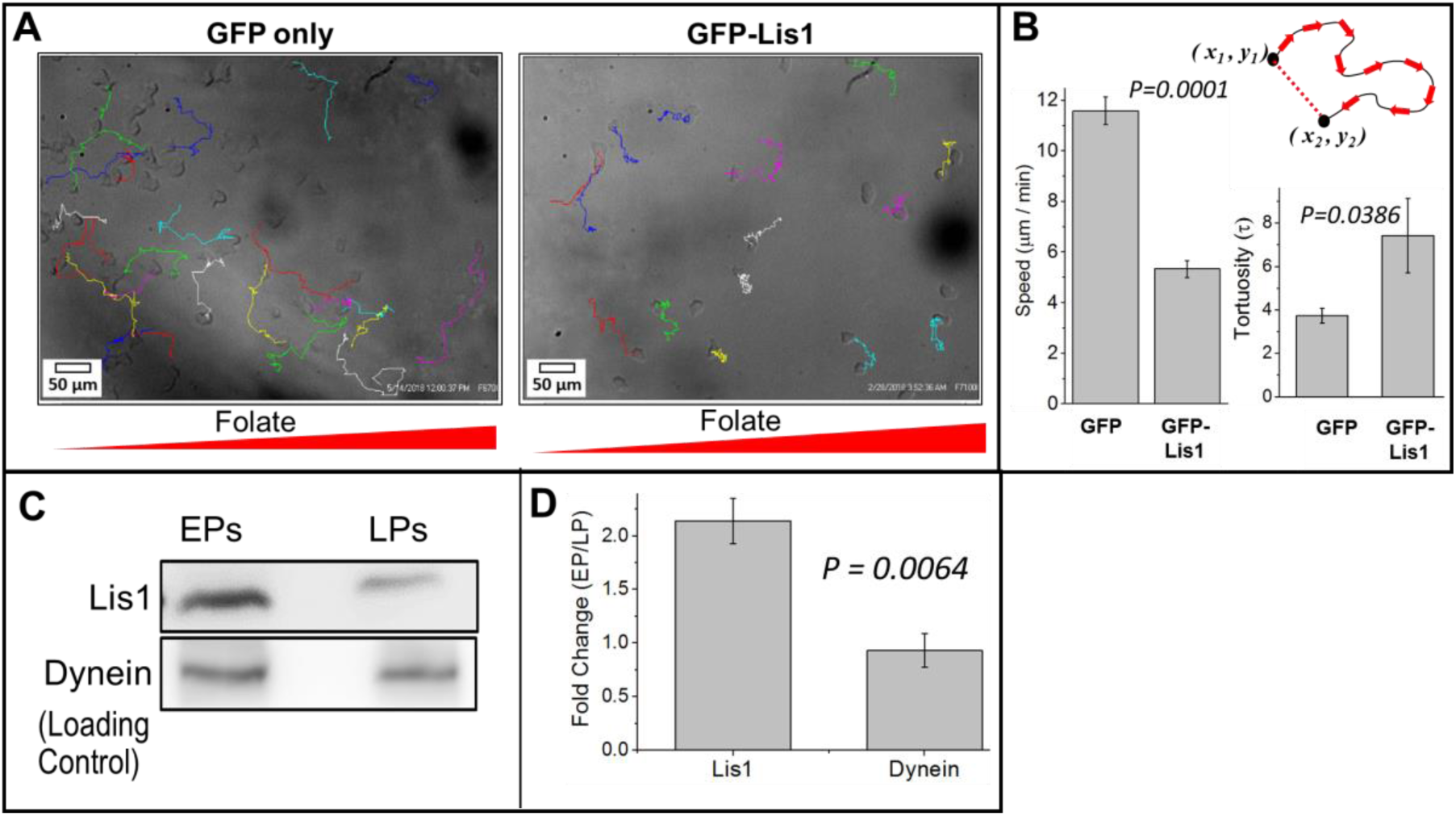
Lis1 overexpression causes cell motility defects. Lis1 is enriched on Early phagosomes (EPs). A. DIC images of Dictyostelium (AX2) cells overexpressing GFP only or GFP-Lis1 constructs, overlaid with migration traces recorded over 40mins. B. Speed and Tortuosity measurements of migration. Schematic describes the meaning of tortuosity (see main text). Lis1 overexpression leads to reduction in speed and increase in tortuosity of migration. n = 30 cells in each set across 3 independent experiments. Error bars are SEM. C. Representative immuno-blots showing amounts of Lis1 and Dynein on early phagosomes (EPs) or late phagosomes (LPs). Total protein extracted from equal number of EPs and LPs was loaded in each lane. Dynein is a loading control (see main text). Lis1 antiserum was generated and used for this experiment (Methods and Supp Fig 2). D. Densitometric quantification of experiment in D. Lis1 is ~2 fold higher on EPs while Dynein is equal. Error bars indicated SEM.

The denominator is the square root of [(x_2_−x_1_)^2^−(y_2_−y_1_)^2^]. For control cells we calculated τ = 3.73±0.33 from 30 cells across 3 independent experiments. However, GFP-Lis1 cells displayed a loss of directionality with highly tortuous path (τ = 7.42±1.7; Fig 5B). Thus, overexpression of Lis1 caused a defect in cell migration of *Dictyostelium*, which is a process dependent on actin dynamics [21][22]. Our result is not consistent with findings in reference [27], where expression of Lis1-GFP leads to an increase in distance migrated by neuronal cell re-aggregates. However our results agree with studies in [28], where radial migration of neurons in developing mouse brain was delayed upon expression of full length Lis1. As mentioned earlier, it is possible that Lis1 affects actin dynamics locally (e.g. within pseudopods during cell migration), and therefore varying Lis1 (e.g. by knockdown or overexpression) causes no visible difference in the F/G actin ratio.

### Actin and Lis1 are enriched on Early Phagosomes as compared to Late Phagosomes

We have reported significant changes in the lipid composition of a phagosome membrane as it matures from an “early phagosome” (EP) to “late phagosome” (LP). Specifically, a dramatic increase in cholesterol[18] and ceramide[29] was observed on LPs. Dynein motors were shown to cluster into cholesterol-rich microdomains (lipid rafts) on LPs [18]. This clustering enhanced the transport of LPs towards lysosomes that is driven cooperatively by multiple dynein motors within a cluster. A glycolipid from the pathogenic parasite *Leishmania Donovani* (causes Leishmaniasis) disrupted this clustering to inhibit the motion of LPs. If the lipid membrane changes, it is also possible that some proteins (e.g. Lis1 and actin) change on phagosomes with maturation. Our data suggests a specific role for Lis1 in actin cup formation at the early stage of phagocytosis. If this is true, Lis1 levels may be higher on EPs as compared to LPs. Indeed, western blotting of purified EPs and LPs revealed that Lis1 was ~2-fold elevated on EPs (Fig 5C, 5D). We have previously reported that dynein levels are unchanged across the EP-to-LP transition [18]. Dynein was therefore used as a loading control in Fig 5C. We do not know why Lis1 levels are reduced on late phagosomes, however changes in the proteome of the phagosome with maturation are known [30].

## Discussion

We have shown that both knockdown and overexpression of Lis1 cause defects in phagocytosis. This is likely due to perturbed actin dynamics because membrane protrusions around particles are incomplete upon Lis1 perturbation. The inhibitory effect of Lis1 overexpression on phagocytosis was intriguing, and can be explained in two ways. First, overexpression may cause a dominant-negative effect if the overexpressed Lis1 protein is non-functional. This is because Lis1 functions as a dimer in cells, and endogenous Lis1 may get sequestered into inactive dimers by overexpressed non-functional Lis1. A dominant-negative form of Lis1 results in abnormal microtubule morphology and increased nucleus-to-centrosome distance[11]. However, our data (Figs 2A, 2C) does not favour an explanation of the results in terms of a dominant-negative effect of Lis1. A second possibility is that the over expressed Lis1 is active, and recruits excess actin nucleating machinery locally to the site of phagocytosis, thereby rendering the actin cup less dynamic and inhibiting phagocytosis. We favour this possibility because stabilizing F-actin by drug treatment also inhibited phagocytosis (Fig 2C, 2D). Notably, a phenomenon of cortical actin balance has been reported using atomic force microscopy and computer simulations [20]. The authors report that an optimal cortical tension is necessary for efficient mitosis of HeLa cells. Both reducing and increasing cortical tension from its endogenous levels leads to perturbation in cell division.

Lis1 could be acting as a scaffolding protein to recruit components of the actin nucleation machinery at the site of phagocytosis. Indeed, Lis1 interacts with Rac1 and IQGAP [5], both of which localize to the phagocytic cup[31][32]. Lis1 is involved in regulating extra-cellular adhesion complexes for cell-cell[33] or cell-substratum adhesion during cell migration [6], where the cortical machinery mediates translocation of the cell membrane against a rigid substratum. In a similar fashion, phagocytosis also involves activity of the cortical machinery to deform the cell membrane and exert force on the phagocytosed particle to drive its inward movement[34]. The biochemical pathways governing phagocytosis and cell motility seem to be largely conserved[15][4][9], suggesting similarities in the underlying molecular machinery that drives both these processes.

It is known that Lis1 induces a persistent force-producing state for Dynein [35]. We found that dynein-teams on LPs generate persistent force against an optical trap, but this persistence was reduced for EPs [18]. It is unlikely that the persistent force on LPs is caused by an effect of Lis1 on dynein because Lis1 levels are lower on LPs compared to EPs (FIG 5C). Rather, clustering of dynein on a cholesterol rich platform on the LPs likely causes persistent force generation by many dyneins. This is supported by the observation that a glycolipid purified from the pathogen *Leishmania donovani* disrupts the cholesterol rich platforms and also the persistent force generation by dynein on LPs [18]. In conclusion, we have demonstrated a role of Lis1 in phagocytosis. However, we do not have structural insight of how Lis1 scaffolds the actin nucleation machinery. While a role for Lis1 in modulating dynein function is well accepted [20], less is known about Lis1 function in controlling the actin machinery. Targeted mutagenesis in Lis1 coupled with the phagocytosis assay described here may help identify Lis1 mutations relevant to phagocytosis. A good way to start is to use *pafah1b1* mutations in Lissencephaly patients[36][37]. Screening point mutants of Lis1 for phagocytosis and migration defects might also help in identifying the domain of Lis1 which scaffolds the actin nucleation machinery.

## Materials and Methods

### Plasmids and cloning

Mammalian Lis1 shRNA construct was a kind gift from Prof. Subba Rao Gangi Shetty (IISc, Bangalore) and mammalian Lis1-GFP construct was a gift from Prof. Deanna Smith (University of South Carolina). For cloning Dictyostelium Lis1 (Dd Lis1), total RNA was isolated from Dictyostelium AX2 cells in exponential phase. cDNA was prepared from total RNA and Lis1 cDNA was PCR amplified (Forward primer: G**TCTAGA**TTATTGTAATTTCCAAAC, Reverse primer: GT**GGATCC**ATGGTATTAACTTCA). Lis1 was cloned after the C terminus of GFP tag in pTX-GFP vector between BamHI and XbaI sites.

### Antibodies

Mammalian Lis1(ab2607), Kinesin-2(ab11259) antibodies were obtained from Abcam. GAPDH antibody was obtained from AbClonal Inc. The antibodies were used at 1:3000, 1:2000 and 1:2000 working dilutions respectively.

The source and protocol for generation of Dicty Dynein antibody has been described in detail[38]. For Dicty Lis1 antibody, DdLis1 was cloned in the bacterial vector pET28a between NdeI and HindIII sites. Purification of Dd Lis1 protein was done by expressing this construct in BL21 cells followed by affinity chromatography using Ni-NTA beads. Final dialysis of purified protein was performed in PBS. Bacterially purified Dd Lis1 was used as the antigen for Lis1 antibody generation in mice. 150 µg purified Dd Lis1 protein (100µl) was resuspended in 100ul of Freunds complete adjuvant and was injected intra-peritoneally in each of the two Swiss mice. Pre-immune serum was collected from these animals before injection of antigen. Primary injection was followed by three booster injections of 100µg Dd Lis1 in incomplete adjuvant at regular intervals of one month. After one month of the third booster, animals were sacrificed to obtain blood. Serum was separated from total blood by letting it stand at room temperature for 2 hours followed by a low speed spin at 3000rpm at 4 °C. The clarified serum obtained as supernatant was used as Lis1 antibody. 1 in 2000 dilution of serum in milk was used for detecting Lis1 band by western blotting. Lis1 serum identified Lis1 protein at the correct size from Dictyostelium lysates and purified protein whereas no band was obtained with control pre-immune serum. Dynein antiserum was also used as 1 in 2000 diluted in milk.

### Cell culture

*Dictyostelium:* AX2 cells were maintained in HL5 media (Formedium, UK) on 10cm cell culture dishes containing 100 µg/mL Penicillin/Streptomycin, at 22°C. Cells were induced for development by plating them with *Klebsiella aerogens* on SM/5 agar plates. Developed spore bodies were picked and frozen in phosphate buffered glycerol as aliquots and stored at −80°C for long-term storage.

RAW 264.7 Macrophages: Mouse macrophage cells (hereafter RAW cells) were cultured in DMEM-High glucose (Sigma) supplied with 10% (v/v) heat inactivated foetal bovine serum (Himedia), under standard culture conditions (37°C, 5% CO_2_, Humidified incubator.)

### Cell Transduction

Lis1 shRNA Lentiviral particles were prepared in HEK293T cells and RAW 264.7 macrophages were transduced with these particles using protocol as recommended by Addgene. Transductant cells were selected on Puromycin (2.5 µg/mL). Stocks were frozen in liquid nitrogen. Lis1 knockdown was validated by Western Blotting. Lis1 band was identified at correct size in scrambled shRNA cells, but not in Lis1 shRNA cells.

### Transfection

#### Dictyostelium

AX2 cells were electroporated with either empty pTX-GFP or pTX-GFP-Lis1 plasmid in 0.1cm gap electrode cuvettes using square wave protocol of BioRad Gene Pulser XCell (2 pulses each of 650V, 1 ms duration, 1500 ms interval). Transformed cells were cultured on HL5 medium containing 10µg/mL G418 as selectable resistance marker, sub-cultured from clonal colonies at 22°C and stored at −80°C as mentioned before.

#### RAW 264.7 Macrophages

RAW cells were electroporated as described in [12]. Briefly, 7.5 × 10^6^ cells were electroporated with plasmids of interest in 0.4cm gap electrode cuvettes using the exponential protocol of BioRad Gene Pulser XCell (250V, 960μF capacitance). Transfected cells were washed once in ice-cold PBS (pH 7.4) by centrifugation (200g for 5mins, 4°C) and plated on coverslips (CORNING no. 1.5, 22mm X 22mm) overnight before experiment.

### Phagocytic Index Assay

#### Dictyostelium

750nm diameter latex beads (PolySciences Inc.) were washed thrice with Sorenson’s phosphate buffer and sonicated briefly. Cells were scraped from cultured dishes, washed with Sorenson’s buffer twice, and counted in hemocytometer. Cell density was adjusted to 6 × 10^6^ /mL by diluting the suspension in HL5 medium. Cells were temporarily stored as 50μL suspension aliquots at 22°C. LatrunculinA, Jasplakinolide were diluted in DMSO at stock concentrations of 1mM each. All the reagents were obtained from Sigma. For initiating phagocytosis, a diluted bead suspension was directly added to one aliquot of cells, at 100:1 cell to bead ratio. Specified drug was diluted appropriately during experiment. For control experiment, equal volume of DMSO was added as vehicle. Cells were thoroughly mixed with beads by gentle tapping, and phagocytosis was allowed to progress for 5mins. Cells were flattened on coverslips (CORNING no.1.5, 40mm X 22mm) using 0.8% phosphate agar pieces (1cm X 1cm). Excess bead solution was gently aspirated out using wicks. Snapshots of individual cells were acquired on NIKON TE2000 DIC microscope, 100X oil objective N.A. 1.4, with Cohu 4910 camera. Phagocytic Index was defined as number of beads phagocytosed per cell.

#### RAW macrophages

2.5µm diameter silica beads (PolySciences Inc.) were washed thrice with DMEM (Sigma) by centrifugation (200g for 5mins) and briefly sonicated. Transfected (or control) RAW cells cultured overnight on coverslips, as mentioned before, and were washed once with PBS, and re-supplied with culture medium. Phagocytosis was initiated by directly adding the bead suspension to coverslips. Cells were incubated in humidified incubator at 37°C and 5% CO_2_ for 5 mins. Post bead pulse, cells were washed twice with ice cold PBS to remove non-phagocytosed beads, and fixed in 4% buffered paraformaldehyde (PFA), and further processed for fluorescence staining protocol.

Staining for fluorescence microscopy was performed by permeabilizing cells in PBS containing 5% v/v Triton X-100 (PBS-T), following blocking in 5% (w/v) BSA solution prepared in PBS-T (Abdil). Post blocking, cells were stained with Rhodamine-phalloidin at 1:400 dilution in Abdil for 15mins at room temperature. Coverslips were mounted on microscopy slides in VectaShield mounting medium (VectorLabs). Images were acquired on Olympus FV1200 CLSM under 100X Oil objective (N.A. 1.4). Entire cell volumes of individual cells were acquired as Z-stacks. Image analysis was performed by manually counting beads per cell in ImageJ. Beads appeared as hollow spheres on green (GFP) background.

### Measurement of Phagocytic Cup

*Dictyostelium* cells cultured on coverslips were challenged with 2.5μm diameter silica beads as described before, with a minor modification. Instead of 5min pulse, the cells were incubated with beads for only 2.5 mins to let phagocytosis complete partially. The cells were stained, mounted and imaged as described before. The imaging conditions were kept consistent across samples and experiment sets. Arc-like rhodamine-positive membrane protrusion surrounding the beads was chosen as ROI for analysis. The length of cup (arc of a circle) was calculated using polar transformer plugin of ImageJ (See Results).

### Preparation of cell lysate and western blotting

Dictyostelium cells either expressing pTX-GFP (empty vector control) or pTX-GFP-Lis1 (Test) were scraped from 10cm confluent dishes, washed once by centrifugation, and resuspended in phosphate buffered 0.1% Triton X-100 solution, to obtain total cell lysates.

RAW cells expressing either expressing scrambled or Lis1 shRNA were scraped from confluent T25 flasks, washed once in PBS by centrifugation, and resuspended in 0.1% Triton X-100 solution of phosphate buffered saline, to obtain total cell lysate.

For immunoblotting, the blot was blocked in 5% (w/v) non-fat dry milk prepared in Tris-buffered 0.1% (v/v) Tween-20 (TBS-T) for 2 hours. It was then probed with primary antibodies made in 5% (w/v) TBS-T solution of BSA (Abdil) for 1.5 hours. After thorough washing with TBS-T, Horse-radish peroxidase conjugated donkey secondary antibody (Sigma; made in 5% TBS-T solution of milk) was added to the blot (1:10000 dilution) and incubated for 1.5 hours. All the incubations were carried out at room temperature. Blots were developed using chemi-luminescence substrate (Millipore, Merck) on AI600 blot developer (GE).

### Pulldown of HisGFP Lis1 from *Dictyostelium* cell lysate

1 mg of HisGFP and HisGFPLis1 cell lysates were incubated with nickel beads, previously blocked with 5% milk for 1 hour. Incubation of HisGFP and HisGFP Lis1 lysates with blocked beads was carried out at 4 degrees for 3 hours for His pull down. Beads were then extensively washed three times in Tris buffer pH 8 containing 1M NaCl. His tagged complexes were finally eluted from beads by adding Tris buffer pH 8 containing 400mM imidazole.

### Cell Migration assay

Plates were prepared by cutting out Agarose troughs from 1% phosphate agar. One trough was filled with buffered 2mM folate solution (pH 7.4). The plate was incubated at 22°C for 10 mins to let the folate form chemotactic gradient in the Agarose. For the experiment, approximately 10^4^ cells either expressing pTX-GFP or pTX-GFP-Lis1 were added to empty trough. The cell motion was then recorded using time lapse microscopy using NIKON TE2000 microscope (10X objective) for 40 minutes. Cells were tracked using manual tracker plugin of ImageJ. Migration speed and persistence (tortuosity) was calculated for each cell type using pair-wise distance formula of each cell’s co-ordinate obtained.

### Super-Resolution Microscopy (dSTORM)

For dSTORM imaging, RAW cells were cultured on coverslips expressing either Lis1 shRNA or scrambled shRNA. Cells were incubated with 2.5µm diameter silica beads for 2.5 minutes at 37°C. Cells were then fixed and incubated with Rabbit anti-Lis1 antibody (Abcam) at 1:300 dilution, and further stained with Alexa555-conjugated anti-rabbit secondary antibody (Invitrogen) at 1:500 dilution. Cells were counterstained with Alexa488-phalloidin (Invitrogen) at 1:250 dilution. Samples were mounted in STORM buffer as specified by the manufacturer.

Data was acquired on the NanoImager S Mark II from ONI (Oxford NanoImaging) with the lasers 405nm/150mW, 473nm/1W, 561nm/1W, 640nm/1W and dual emission channels split at 560nm. 12500 frames were acquired sequentially at 100% laser power in both samples for each channel. Image reconstruction and post-acquisition processing for background filtering was done on NimOS (Version 1) from ONI. For analysis, a line was drawn radially from the bead centre into the cell, and line intensities were plotted for both channels using ImageJ.

## ACKNOWLEDGEMENTS

Mammalian Lis1 shRNA construct was a kind gift from Prof. Subba Rao Gangi Shetty (IISc, Bangalore). Mammalian Lis1-GFP construct was a gift from Prof. Deanna Smith (University of South Carolina). We thank Dr. Pradeep Barak and Ashwin D’Souza for technical help with experiments and for discussions. We also thank Dr. A. Ghosh for comments on the manuscript. RM acknowledges funding through an International Senior Research Fellowship from the Wellcome Trust UK (grant WT079214MA).

---

**Supplementary Figure 1:**
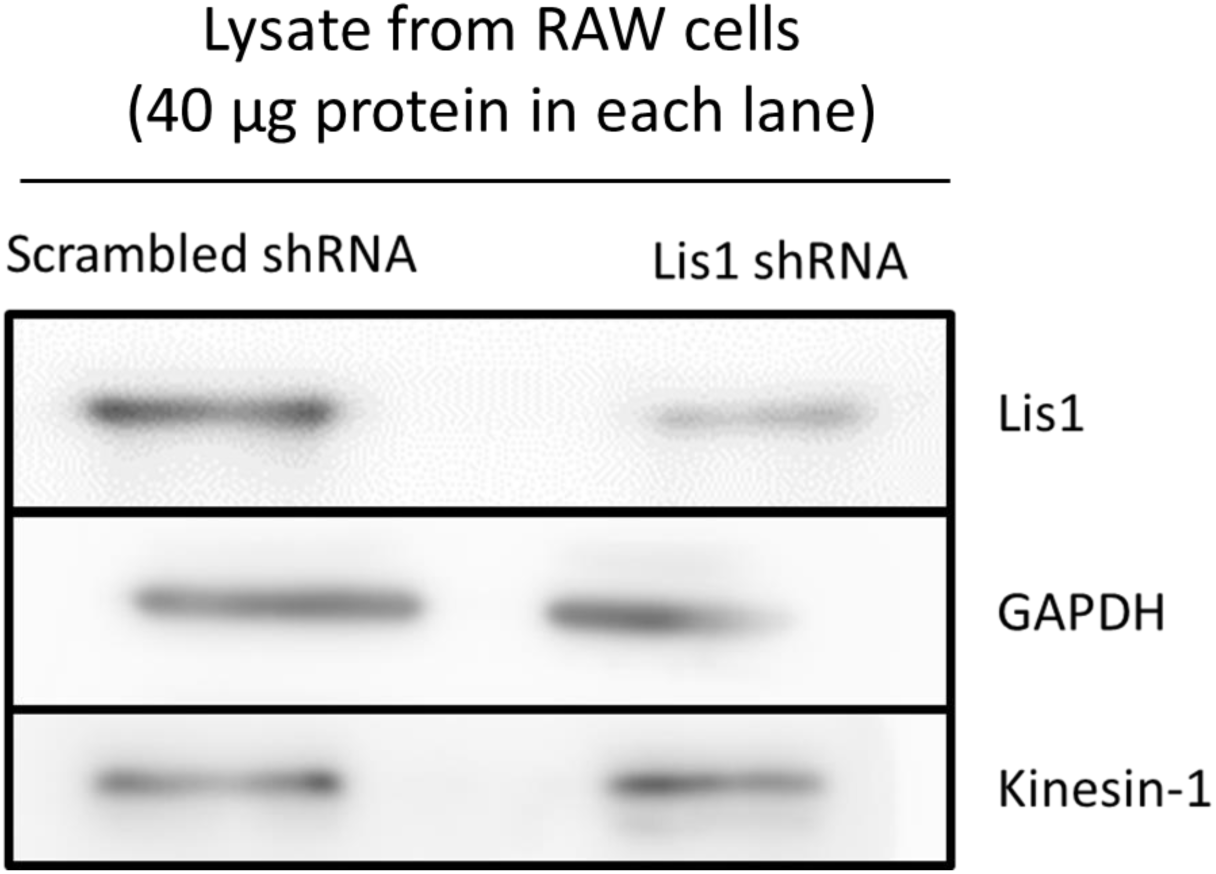
Validating Lis1 knockdown in RAW cell lysate. Western Blot showing specific knockdown of Lis1 in RAW cell lysate (40 µg) upon expression of *lis1* shRNA as compared to scrambled control using viral transduction. GAPDH and Kinesin-1 are used as loading controls. Lis1 antiserum was generated and used for this experiment (Methods and Supp Fig 2).

**Supplementary Figure 2:**
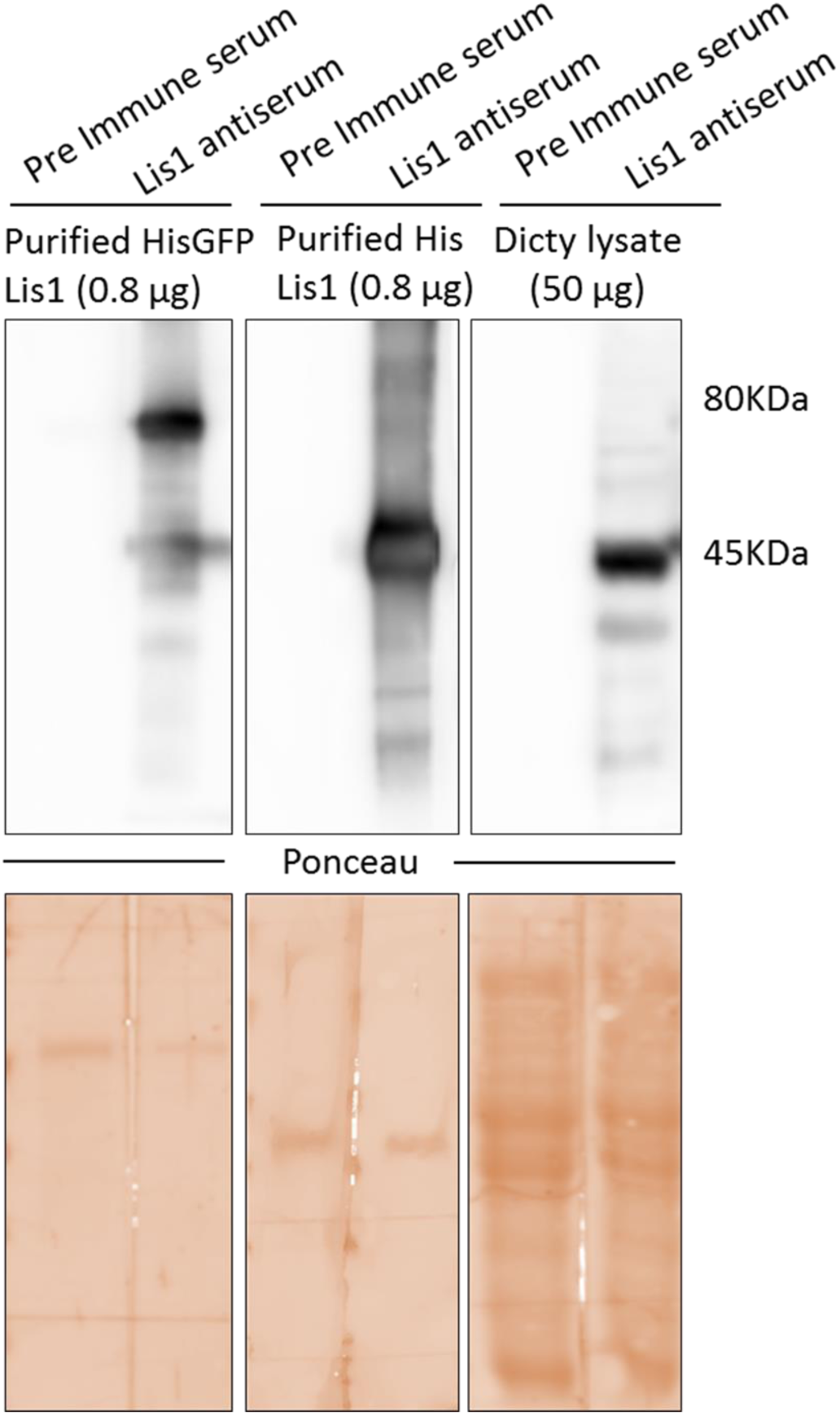
Validating Lis1 antiserum using western blotting. Western blot showing the specificity of Lis1 antiserum as compared to pre immune serum. Both pre immune and Lis1 antiserum were tested on 0.8 µg of purified HisGFPLis1 protein (purified from *Dictyostelium*), 0.8 µg of HisLis1 protein (purified from bacteria) as well as on 50µg of *Dictyostelium* wild type crude lysate. Specific bands corresponding to size of HisGFPLis1 (80KDa) and Lis1 (45KDa) were obtained when probed with Lis1 antiserum. No signal was obtained on blots incubated with pre immune serum even at high exposure times.

